# *Leucaena* and *Acaia* leaf litter negatively affect the survivorship of tadpoles of the South Asian frog *Microhyla ornata*

**DOI:** 10.1101/2024.01.22.576772

**Authors:** Aranya Pathak-Broome, Vivek Gour-Broome, Shomen Mukherjee

## Abstract

Leaf litter from terrestrial plants forms an essential source of organic food matter for many freshwater aquatic organisms. However, leaves of some species are known to alter water quality, affecting the development and survivorship of amphibians. While amphibians from North and South America are affected by terrestrial leaf litter, similar studies are missing from Asia, a region with high amphibian diversity (mainly from the south and south-east Asia). At different tadpole densities, we tested the effect of different leaf litter (two non-native trees and a grass species) on the survivorship of ornate narrow-mouthed frog tadpoles (*Microhyla ornata*), a common frog in South Asia. We found the tadpole probability of survival extremely low in *Leucaena* but high in *Themeda*. While the odds of survivorship were nearly twelvefold higher in *Acacia* than *Leucaena*, it was not as high as in *Themeda*. A tadpole also had a lower odds of survival if raised in a high-density environment. In addition, *Leucaena* litter water had significantly higher pH levels than other litter types. Since both *Leucaena* and *Acacia* are non-native trees actively grown for agroforestry in South Asia, our results highlight another potential threat to amphibians in this region. Given the high amphibian diversity in this region, there is an urgent need for similar studies on other anurans and aquatic organisms.

---

Terrestrial leaf litter is an important source of carbon for many aquatic systems (Brinson et al., 1981; Fisher & Likens, 1973; Polis et al., 1997; Vannote et al., 1980). These dead plant materials fuel the aquatic food webs (Gessner et al., 1999), thus affecting the community composition, diversity, and growth of aquatic organisms (Polis et al., 1997). Litter from different species of terrestrial plants can affect aquatic organisms in wetlands differently. Hence it is vital to understand species-specific effects (Stoler and Relyea 2011). Since the spread of non-native terrestrial plants is a cause of concern for many native fauna (Blossey, 1999), non-native litter subsidies may affect some aquatic fauna negatively since they have not evolved with them (Brown et al., 2006).

Water-soluble phytochemicals produced by invasive plants can have numerous direct and indirect effects on native organisms and affect the survivorship of anuran tadpoles (Brown et al., 2006; Watling et al., 2011). Depending on the species of plant leaf litter present, changes in aquatic environments could be due to plant secondary compounds or differential algal growth, bacterial production, and phytoplankton biomass (Rubbo et al., 2008). Due to varying litter decomposition rates and allelopathic effects, non-native plant litter can inhibit bacterial, phytoplankton, and periphyton growth, affecting the abundance of higher trophic-level organisms (Maerz et al., 2005). The adverse effects of non-native leaf litter, in the form of low development rates and survivorship of tadpoles, are well documented for North American species of anurans (Brown et al., 2006).

Ephemeral ponds (e.g., temporary ponds) are essential for developing larval stages of several anuran species and invertebrates. Compared to flowing water bodies such as streams and rivers, these small seasonal pools may be more affected because of their small size, thus leading to a higher concentration of tannins and other water-soluble phytochemicals. Also, since litter affects the algal composition and hence has a bottom-up effect on the tadpoles (Brown et al., 2006), it is essential to understand the impact of litter subsidy on aquatic organisms in smaller water bodies.

Almost all the studies on the effect of terrestrial leaf litter on anuran reproduction, growth, and survivorship are from the Americas. To our knowledge, we are unaware of any such study from Asia. Such a study is urgently needed since amphibian diversity is relatively high in Asia (particularly in South and Southeast Asia), and because populations are highly threatened due to habitat loss (Sodhi et al., 2004, Rowley et al., 2010). Also, like most other continents, this region has many non-native and invasive species of plants.

In Western India (Pune, Maharashtra), we experimented to understand the effects of two non-native trees (*Acacia auriculiformis* and *Leucaena leucacephala*) and a native grass (*Themeda quadrivalvis*) litter on the survivorship of *Microhyla ornata* tadpoles and water pH. These plants contribute leaf material to temporary ponds in many parts of India. We further tested if the density of tadpoles (crossed with leaf litter) influenced their survivorship.

## MATERIALS AND METHODS

### Study species

*Microhyla ornata* (family Microhylidae), the ornate narrow-mouthed frog, is a common nocturnal frog distributed across Bangladesh, Bhutan, India, Myanmar, Nepal, Pakistan, and Sri Lanka. The tadpoles of this species feed on algal and detrital material. They breed in the temporary ponds of scrub forests and grassland habitats (Dutta et al., 2008).

*Acacia auriculiformis* (*Acacia*) is a tree native to Papua New Guinea and Australia and has been introduced to tropical and sub-tropical areas as an ornamental tree and for reforestation. In India, Acacia was introduced in 1946, and used as ornamental plant, but later for fuel wood and restoration of degraded land (Turnbull et al., 1998). *Leucaena leucocephala* (*Leucaena;* common name: Subabul) is a tree native to southern Mexico and northern Central America that has naturalized throughout the tropics. *Leucaena* was introduced to India in the 1980s as a dry-land forest crop and is a good nitrogen-fixing tree and fodder crop. *Themeda quadrivalvis* (*Themeda*) is a species of annual grass native to India.

### Experimental design

We conducted an outdoor experiment using food containers at Jambe Environmental Farm, Pune (600m above sea level, 18°37’ N, 73°43’ E). Each food container (1L) contained 900 ml of rainwater and 2 grams of dried leaf litter. Although standard mesocosm experiments use 1kg litter/1000L of water, our amount was higher since many non-native plantations are monocultures. *Acacia, Leucaena*, and *Themeda* leaf litter were collected after senescence, washed in rainwater, and dried before use. Although we collected tadpoles from natural ponds that did not have *Acacia* or *Leucaena* trees next to them, these plants commonly occur near ponds in the study area and other parts of India.

One week before the start of the experiment (7 July 2018), leaf litter and rainwater were added to the containers to allow the natural colonization of algae. On 12 July 2018, we collected *Microhyla ornata* tadpoles (Gosner stage 23) from nearby temporary ponds formed during monsoon rain. They were from multiple clutches but of roughly the same size and the same Gosner stage. Tadpoles were added to the containers on 14 July 2018 (start date), and the experiment lasted a month.

Thirty-six containers were set up in a block design. Each block consisted of six-leaf litter treatments: three leaf litter treatments (*A. auriculiformis, L. leucacephala* and *T. quadrivalvis*) crossed with two density treatments (Low – two tadpoles/containers and High – four tadpoles/container). The containers were placed at random within a block, making sure that the pattern was not repeated for multiple blocks. The blocks were all kept under a tin roof to prevent rainwater from falling in them, but they received sunlight and other natural conditions from the sides. The water level in the containers was allowed to drop naturally, thus replicating the conditions of a temporary pond.

Survivorship for each tadpole was converted into binomial data and subsequently used as a response variable in a binomial logistic regression in R [“glm(model, family=binomial)”; Version 4.0.3, R 2021]. Next, we measured the odds of survival in the different *leaf litter* and *density* treatments (the two predictor variables). The regression results were interpreted after converting the log-odds ratio to odds ratio. To test for overall statistical significance of the logistic regression, McFadden’s Pseudo *R*^2^ value (the overall effect size) was also calculated to estimate the goodness-of-fit, followed by a chi-square distribution for *P* value. Finally, we calculated the predicted probability of survival (from the logistic regression) when raised in each leaf litter type and across the two density treatments. The binomial logistic regression and following statistics were carried out according to Starmer (2021) in R (Version 4.0.3, R 2021).

We measured the pH of the water in the containers twice, once at the beginning (12 July 2018) and then in the middle (24 July 2018, 17 days after adding litter). A two-way ANOVA was also conducted on the 24^th^ July pH data to test for leaf litter and density effects. We used R function “aov()” (Version 4.0.3, R 2021) for the statistical analysis. We also calculated the Cohen’s *d* and Pearson’s coefficient *r* effect sizes (Cohen, 1988) and categorised the difference as small (Pearson’s *r* > 0.1), medium (Pearson’s *r>0*.*3*) or large difference (Pearson’s *r* > 0.5) based on Cohen (1988) and Sawilowsky (2009).

According to the International Union for Conservation of Nature, *M. ornata* is listed in the category of *Least Concern*. The study fulfilled the requirements of the Institutional Research Board of Azim Premji University, Bangalore, India.

## RESULTS

Tadpole survivorship is significantly affected by both leaf litter type and tadpole density (Binomial regression analysis: McFadden’s Pseudo *R*^2^ = 0.46, *p* <0.001, Table 1). The predicted probability of a tadpole surviving was least in *Leucaena*, intermediate for *Acacia* and highest for *Themeda* litter (Fig. 1). When raised in *Themeda*, tadpoles have 265 and 22.23 times higher chances of survival compared to those raised in *Leucaena* and *Acacia* respectively (Table 1). Similarly, when tadpoles were raised in *Acacia* the odds of surviving significantly increased by a factor of 11.93 compared to *Leucaena* (Table 1). The overall odds of surviving increased by a factor of 3.99 for a tadpole being raised in low density compared to high density (Table 1).

**Table 1:** Binomial regression analysis indicating strong effect of leaf litter and tadpole density on tadpole survivorship.

|  | Estimate | Std. Error | z value | *P* value |
| --- | --- | --- | --- | --- |
| (Intercept) | - 3.05 | 0.72 | - 4.22 | <0.001 |
| Plant (*Acacia*) | 2.48 | 0.73 | 3.39 | <0.001 |
| Plant (*Themeda*) | 5.58 | 1.00 | 5.57 | <0.001 |
| Density (low) | 1.38 | 0.62 | 2.22 | 0.03 |

**Fig.1:**
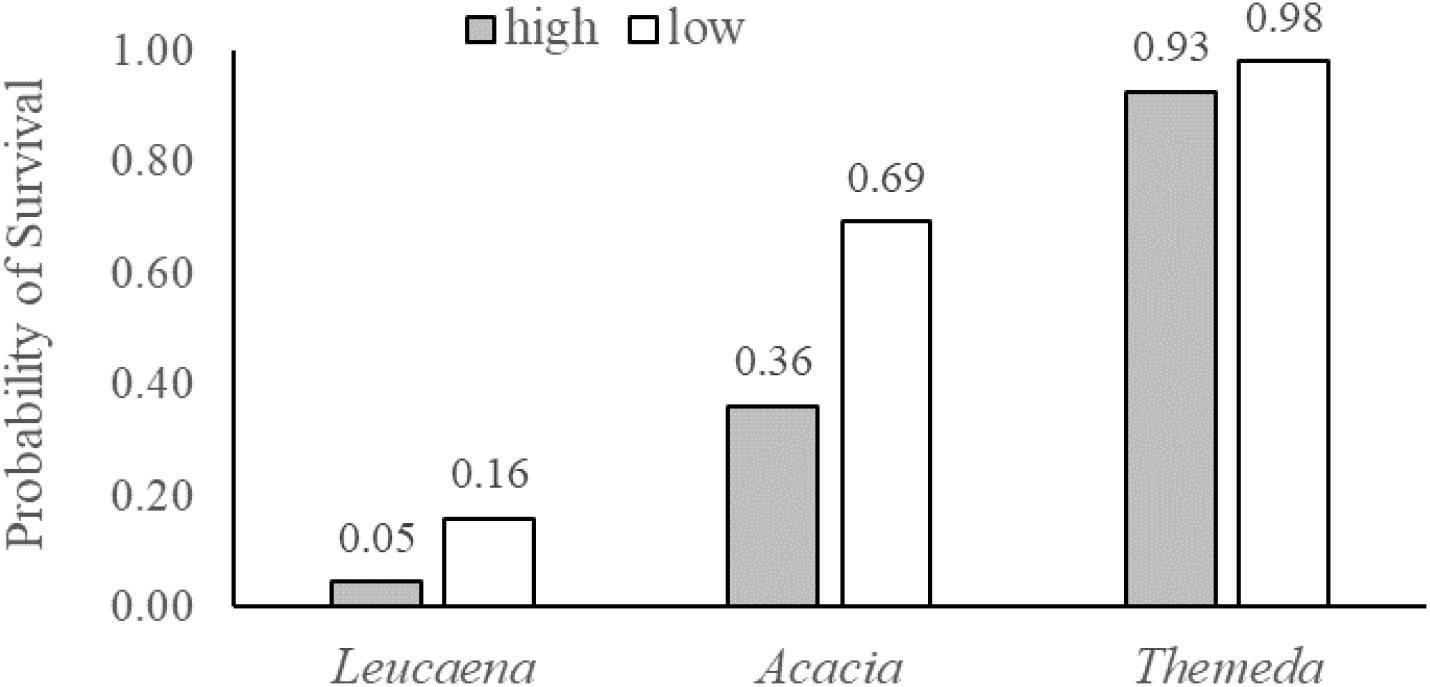
Predicted probability of survival for tadpole across different leaf litter treatment in *high* (grey bar) and *low* (white bar) density. The numbers above the bars are treatment specific predicted probability of survival.

Leaf litter type significantly affect the pH (*F*_2,30_ = 178.34, *p* < 0.001). *Leucaena* litter had the highest pH, followed by *Acacia* and *Themeda*. Tukey post-hoc tests indicated only marginal difference in pH between *Acacia*-*Themeda* (*p* = 0.05; lower in *Themeda*). However, there were strong differences between *Leucaena*-*Themeda* (*p* < 0.001) and *Leucaena*-*Acacia* (*p* < 0.001). Effect size analysis indicated huge difference between *Leucaena* and *Acacia* (Cohen’s *d* = 5.10, *r* = 0.93) and between *Leucaena* and *Themeda* (Cohen’s *d* = 6.11, *r* = 0.95) and a large difference between *Acacia* and *Themeda* (Cohen’s *d* = 1.06, *r* = 0.47). Tadpole density also affected pH (litter type*density; *F*_2,30_ = 4.77, *p = 0*.*02*), however post-hoc tests did not show significant differences between high and low density for any litter treatment (Fig. 2).

**Fig. 2:**
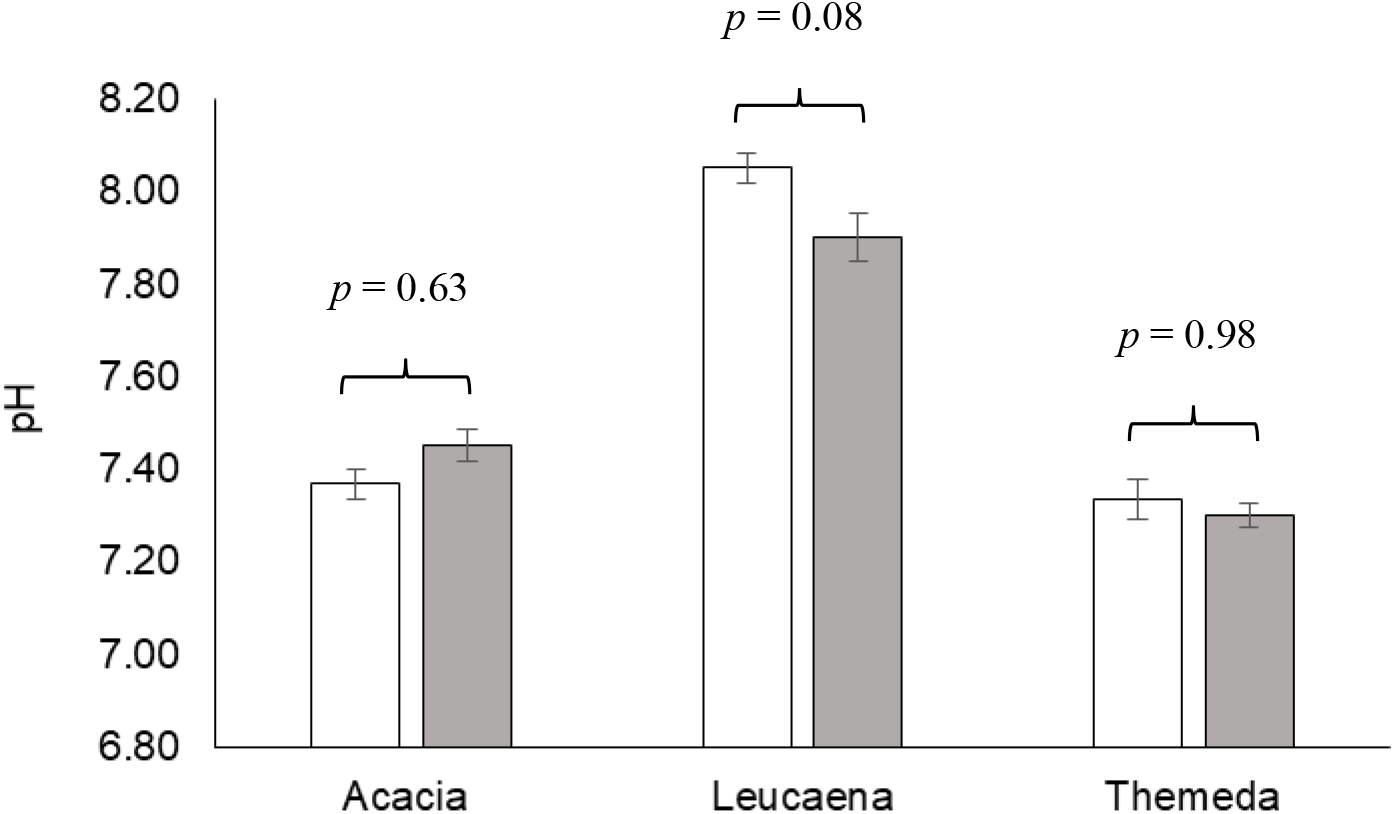
Difference in pH across different leaf litter treatments across *high* (open bar) and *low* (grey bar) density of tadpoles. Error bar indicates Mean ± SE and numbers above the bars indicate post-hoc differences between density treatments.

## DISCUSSION

Our study tried to look at the effect of plant species monocultures on the survivorship of a common frog species. This allowed us to take the first steps towards understanding the relationship between terrestrial plant composition and aquatic fauna in a poorly studied system. Even though the study lacked the ability to discuss the exact mechanisms, it is among the first study of its kind in this biodiverse region that clearly shows how certain types of leaf litter can negatively affect tadpole survivorship. This study is also novel because it tests the effect of two plant species *Leucaena leucacephala* and *Acacia auriculiformis*, that have been introduced in many parts of the tropics but who’s effects on vertebrates (apart from domestic livestock) have not been studied.

What is clear and revealing was the powerful effect of *Leucaena* and the negligible effect of *Themeda* on survivorship (both at low and high density; Fig. 1). One of the possible reasons for such differences in survivorship between the litter types could be the presence of tannins (reactive phenolics). Although we did not measure tannins in our study, *Leucaena* and *Acacia* are known to be allelopathic, and tannins are one of their main allelochemical compounds (Hammond 1995, Oyun 2006). We did not find any study that has reported tannins in *Themeda*. Although some grasses may have tannins in their grain, grass leaves lack tannins (Bernays et al. 1989).

Tannins can affect both the quality and quantity of the food consumed by aquatic organisms such as tadpoles. Gut analysis of American toad tadpoles (*Anaxyrus americanus*) showed differences in algal communities between tadpoles exposed to different litter types (Brown et al., 2006). Since tannins can cause chlorosis in plants (Muthukumar et al., 1985), similar effects are also likely in algae, thus affecting the diet of algae-feeding tadpoles.

Tannins from certain plants at specific concentrations affect aquatic organisms’ development and survival (Earl and Semlitsch 2015). For example, tannin exposure causes gill damage in fish, thus causing high mortality (Temmink *et al*. 1989). It is also one of the leading causes of mortality in amphibians (see Maerz et al., 2005). Although the physiological effects of tannins in amphibians are poorly understood, they affect mammal digestion and growth (see Chung-MacCoubrey et al., 1997; Robbins et al., 1991). Such physiological effects and reduced food availability, in terms of quality and quantity of algae, are the likely reasons for reduced survivorship among the tadpoles. Although we did not quantify algae in our microcosms, future studies could test if tannins from *Leucaena* and *Acacia* cause algal chlorosis and affect algal diversity.

There are likely other mechanisms that could help explain the differences in survivorship. For example, in a detailed and complex mesocosm study, Stoler and Relyea (2015) found leaf litter traits such as soluble carbon and litter decay rate best explained the dynamics of freshwater temporary pond communities (including tadpoles). However, they concluded that the chemical attributes of the leaf litter best explained how ponds would respond to litter, specifically soluble carbon. High amounts of soluble carbon in the water can lower algal growth (via reduced photosynthesis) and increase microbial growth, which in turn decreases the amount of dissolved oxygen, thus negatively affecting the survival of aquatic organisms such as tadpoles (Klug 2002; McIntyre and McCollum 2000; Stoler and Relyea 2015). *Leucaena* likely increased the amount of soluble carbon and triggered a similar chain of events.

*Leucaena* and *Acacia* plantations are common in western India, and their leaf litter contribute organic matter to temporary and permanent freshwater ponds. *Themeda* is a very common species of grass that is native to India. Although the aim of the study was not to specifically compare native vs non-native leaf litter, such a study will be extremely valuable since other studies have found invasive tree leaf litter to affect the survivorship of amphibians. For example, a gut analysis study on the American toad (*Anaxyrus americanus)* tadpoles found different algal communities between those raised on native and non-native litter (Brown et al., 2006). Another study on the same tadpole species found significantly higher mortality in those raised in an invasive plant litter (Amur honeysuckle, *Lonicera maackii*) compared to native plant litter (Watling et al., 2011).

Given the wide distribution of these frogs, there is likely within-species and between-population variation in response to leaf litter. However, our strong effects of *Leucaena* (and *Acacia*) could be because we collected tadpoles from temporary ponds which did not have *Leucaena* trees next to them. Although *Leucaena* is considered a naturalized species, these populations might not have had the time to adapt to these tannins. Future experiments where tadpoles are collected from a wide range of habitats will be extremely valuable for understanding adaption in this species and assessing the conservation threat of these trees.

Although we found significantly higher pH levels (alkaline) in *Leucaena* compared to other litters (Fig. 2), it is unlikely the cause of high tadpole death. Our observed range was well within the range reported for other aquatic producers and consumers (Havas and Rosseland 1995, Stoler and Relyea 2011). Unfortunately, we don’t know if *Leaucaena* litter created more hypoxic conditions, which could explain the high mortality combined with the effect of tannins. Since leachates can have a more substantial effect in non-aerated water (Canhoto et al. 2013), future studies should also measure dissolved oxygen since it can mediate the impact of leachates and, thus, the growth of microbes.

In our study, the mechanisms mentioned above or their combinations could be affecting tadpole survivorship. However, we acknowledge that our experiment cannot pinpoint the exact mechanism (or their interaction) behind the reduced survivorship since we did not quantify leaf litter traits or other abiotic and biotic factors. Future studies should explicitly test these mechanisms across various plant species litter.

Our findings of low survivorship at high density (Fig. 1) are not surprising since there is strong evidence in the literature that intraspecific resource competition affects growth, behaviour and survivorship (Connell, 1961; Light, 1967; Stein et al., 2017). However, what is clear is that the effect of tadpole density is negligible in *Themeda* litter (Fig. 1). The absence of tannins and the presence of organic matter allowed high levels of algal growth, which allowed the tadpoles to thrive even at high density. Future mesocosm studies representing larger aquatic habitats and across a range of leaf litter concentrations and tadpole densities would better help understand this system.

To conclude, there is a clear negative effect of *Leaucaena* on *M. ornata* tadpole survivorship. However, since this is the only study we know from Asia, we urgently need studies that test similar effects (and of other non-native and naturalized tree and shrub species) with other amphibian and plant species.

## Acknowledgements

We thank Leela Gour Broome and the late Ashok Gour Broome (Jambe Environmental Farm, Pune) for permitting the study. We would also like to thank Neema Pathak Broome for help with the experimental set-up, Vijayan Sundararaj for his valuable comments, and Josh Starmer (Statquest) for his help with statistics through his YouTube channel. The study was conducted after the Institutional Research Board approval of Azim Premji University (# 2018-07).

